# A one-pot workflow for spatial proteomics aided by ShapeX

**DOI:** 10.64898/2026.11.24.768900

**Authors:** Rune Daucke, Maike W. Nielsen, Julie B. Jacobsen, Vladyslav V. Tkach, Nadine M. Hammouda, Eva L. Lund, Aušrinė Areškevičiūtė, Bjarne W. Kristensen, Valdemaras Petrosius, Erwin M. Schoof

## Abstract

The rapid growth in instrumental sensitivity has enabled deep proteomic profiling of small tissue regions, increasing the availability and interest in spatial proteomics. However, sample preparation remains a practical bottleneck, as any additional clean-up and transfer steps increase losses of already limited material. Here, we developed a one-pot workflow in which laser-microdissected tissue is collected, de-crosslinked, and digested in the same well, followed by direct injection for liquid chromatography–mass spectrometry. We first evaluated the de-crosslinking and digestion across different tissue inputs. Heating at 95 °C for 15 minutes provided a practical balance between proteome coverage and processing time, reducing the heating period by 75% relative to a 60-minute incubation. Using this condition, we quantified a mean of 2,751, 3,237, 3,786, and 5,046 protein groups from 1,000, 2,000, 5,000, and 10,000 µm² regions, respectively, without match-between-runs. We next evaluated 1,536-well plates for higher-throughput tissue collection and sample preparation. Following transfer for LC–MS analysis, the strongest proteomic signals generally coincided with the intended collection wells, indicating that tissue collection in this format is feasible.

In parallel, we developed ShapeX, a QuPath extension that links tissue annotation to laser microdissection and plate-based sample preparation through customizable plate layouts, annotation tessellation, and coordinated export of sampling and dispensing files. We used ShapeX to map proteomic measurements back to the original tissue regions in two intestinal experiments. Spatial reconstruction recovered opposing gradients in proteins associated with epithelial differentiation and replication and distinguished the sampled glandular epithelium from adjacent stroma. Together, the one-pot preparation and ShapeX provide a practical workflow for image-guided spatial proteomics, from tissue annotation and sample collection to spatial mapping of protein abundance.

## Introduction

Continuous advancements across the field of mass spectrometry–based proteomics have enabled the characterization of the molecular composition of increasingly small amounts of input material [1–3]. This capability has extended to spatial proteomics of formalin-fixed, paraffin-embedded (FFPE) material, with laser capture microdissection (LCM) allowing for the precise collection of specific cellular regions for subsequent molecular characterization. By linking proteomic measurements to tissue morphology, this approach enables the investigation of molecular differences that may be obscured when larger tissue regions are analyzed as a whole [4, 5].

At the same time, advances in computing performance, data storage, and whole-slide imaging have supported the wider adoption of digital pathology, allowing tissue morphology to be examined and annotated in a digital environment [6]. Automated image analysis and artificial intelligence (AI) further expand the scale at which tissue structures and cellular populations can be identified and classified [7, 8]. These developments enable increasingly detailed selection of regions according to their morphology, cellular composition, or pathological features. Realizing the potential of these parallel advances requires both efficient processing of small tissue inputs and a reliable connection between the tissue image, the sampled region, and its collection well.

However, the ability to isolate smaller regions does not by itself ensure that the material can be processed efficiently. As the tissue input decreases, each transfer, clean-up, or additional handling step becomes more consequential [9]. Sample preparation must therefore balance protein extraction and digestion with the need to preserve the limited material available for analysis. Methods such as single-pot, solid-phase-enhanced sample preparation (SP3) [10], filter-aided sample preparation (FASP) [11], and C18 StageTip desalting facilitate contaminant removal and compatibility with LC–MS analysis [12]. Nevertheless, incomplete recovery and adsorption to surfaces during processing can reduce the amount of material ultimately reaching the mass spectrometer.

In Evosep-based proteomics workflows, peptide digests are loaded onto Evotips for desalting and subsequent introduction into the LC–MS system. However, when starting material is limited, even transferring the peptide digest from the collection well into an Evotip can cause sample loss, further reducing the amount available for analysis. These losses can arise from incomplete recovery from the well or peptide adsorption to the surfaces of pipette tips [13]. Minimizing transfers between sample preparation and LC–MS analysis is therefore an important consideration when developing workflows for small tissue inputs [14].

Keeping the sample in its collection well until LC–MS injection offers one way to minimize these transfers. For FFPE tissue, this means carrying out de-crosslinking and proteolytic digestion in the collection well, followed by direct injection of the resulting digest into the LC–MS system. Here, we evaluate this approach as a one-pot workflow for spatial proteomics (Figure 1), connecting tissue collection, sample preparation, and direct aspiration from plates without an intermediate peptide transfer.

**Figure 1.**
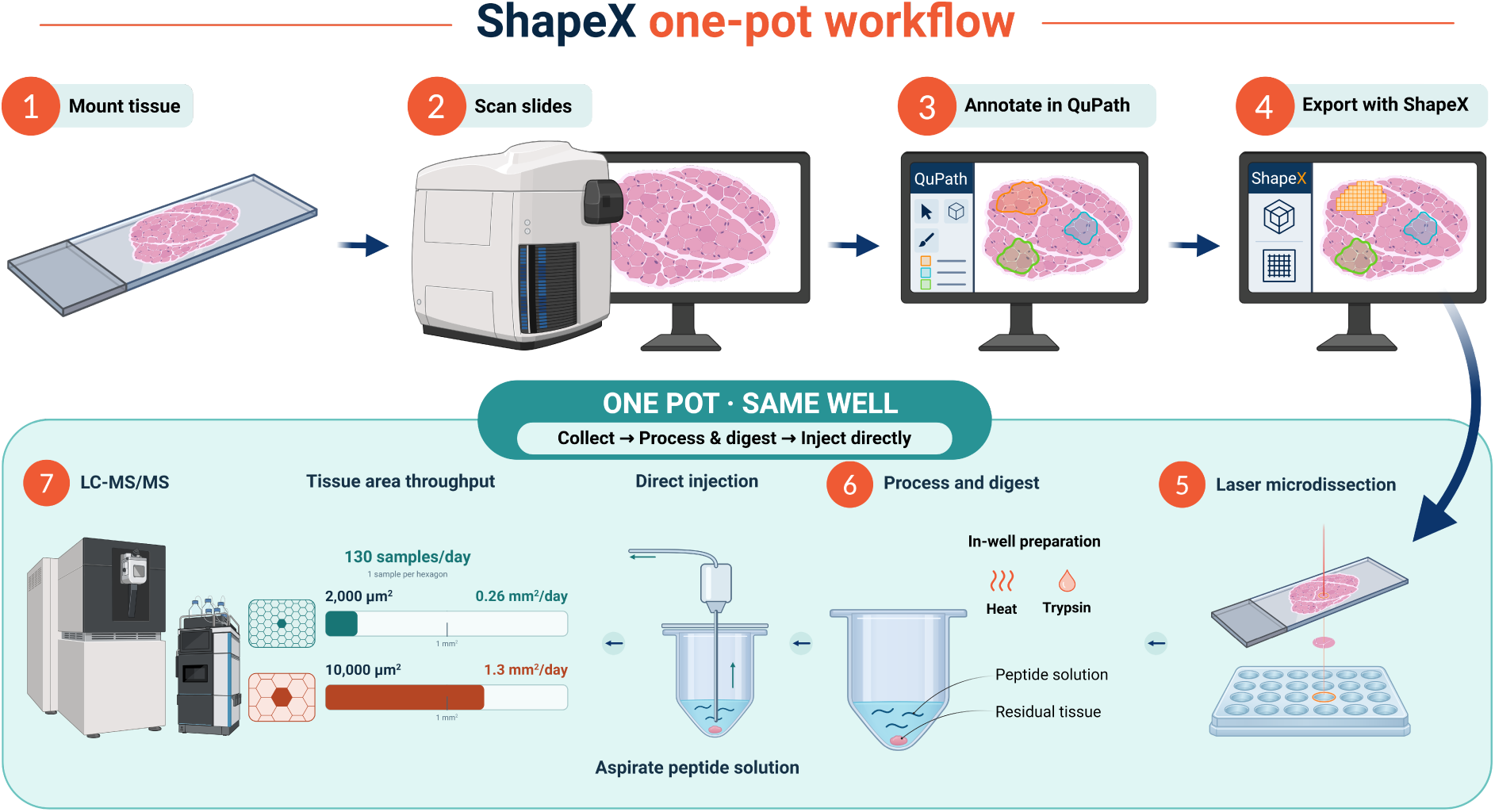
A one-pot workflow for image-guided spatial proteomics. Tissue sections are mounted, scanned, and annotated in QuPath before laser microdissection. ShapeX links the annotations to collection wells and generates files for tissue collection and reagent dispensing. The collected material is heated and digested in the same well, followed by direct aspiration for LC–MS/MS analysis (created using BioRender.com).

Another challenge is translating tissue annotations into practical instructions for sample collection and preparation. Tissue annotations must be linked to destination wells while retaining their original geometry, identity, and spatial relationships. Managing these links can require substantial manual work, particularly when regions are divided into many smaller sampling units, pooled for analysis, or distributed across multiple collection plates. Randomizing sample placement within and across plates to limit the influence of well-position and plate effects adds complexity to collection planning. Connecting pathological knowledge and biological questions with the technical requirements of collection and preparation therefore becomes increasingly important as sampling scales expand.

To address this challenge, we developed ShapeX, an extension for QuPath [15], an established platform for viewing, annotating, and analyzing whole-slide images. ShapeX connects tissue annotations with collection wells and downstream preparation files within the same digital pathology environment. It provides a customizable plate designer, manual or automated allocation of annotations, individual or pooled collection, sample randomization, and negative controls. Its tessellation function divides regions of interest into defined sampling areas. The exported files preserve annotation coordinates and well assignments for microdissection, reagent dispensing, and subsequent spatial reconstruction.

We demonstrate this connection between image-guided sampling and molecular analysis in two intestinal experiments. Using the annotation geometry preserved by ShapeX, we reconstruct protein profiles along sampled structures and map abundance differences between glandular epithelium and adjacent stroma in a separate section.

## Results

### Optimization of sample preparation

From our previous experience with LCM sample preparation [16], tissue debris can remain after heating (de-crosslinking) and trypsin digestion and accumulate during subsequent sample handling. Brightfield microscopy inspection suggested that these steps extract proteins from the microdissected material without necessarily dissolving the entire FFPE fragment. This made us consider whether the peptide-containing solution could be analyzed directly from the collection well, avoiding an additional clean-up and transfer. We therefore developed a workflow in which tissue collection, heating, and digestion are performed in the same well, followed by direct aspiration for LC–MS analysis.

We evaluated heating and trypsin digestion in separate optimization experiments. The primary analysis used searches without match between runs (MBR). Using 5 µm tissue sections, we compared heating at 70 and 95 °C for 5, 15, and 60 minutes across 1,000, 2,000, 5,000, and 10,000 µm² tissue inputs, with matched zero-area controls (Figure 2A). Quality control excluded 36 of 300 heating measurements. After 5 minutes at 95 °C, mean coverage ranged from 2,320 to 4,621 protein groups across the tissue inputs, representing increases of 24–36% over 5 minutes at 70 °C. Extending the 95 °C incubation to 15 minutes increased mean coverage to 2,751–5,046 protein groups, 20–46% higher than the corresponding 70 °C conditions. Further extension from 15 to 60 minutes at 95 °C provided approximately 1–5% additional coverage at most inputs, with the largest gain at 5,000 µm², where coverage increased from 3,786 to 4,316 protein groups (14%). Median abundance coefficients of variation (CVs) were typically around 20–30% and tended to be higher at lower tissue inputs. The 2,000 µm² condition heated at 70 °C for 60 minutes showed unusually low and variable coverage (1,588 ± 1,193 protein groups; mean ± SD), with a median abundance CV of 91% (Figure 2B).

**Figure 2.**
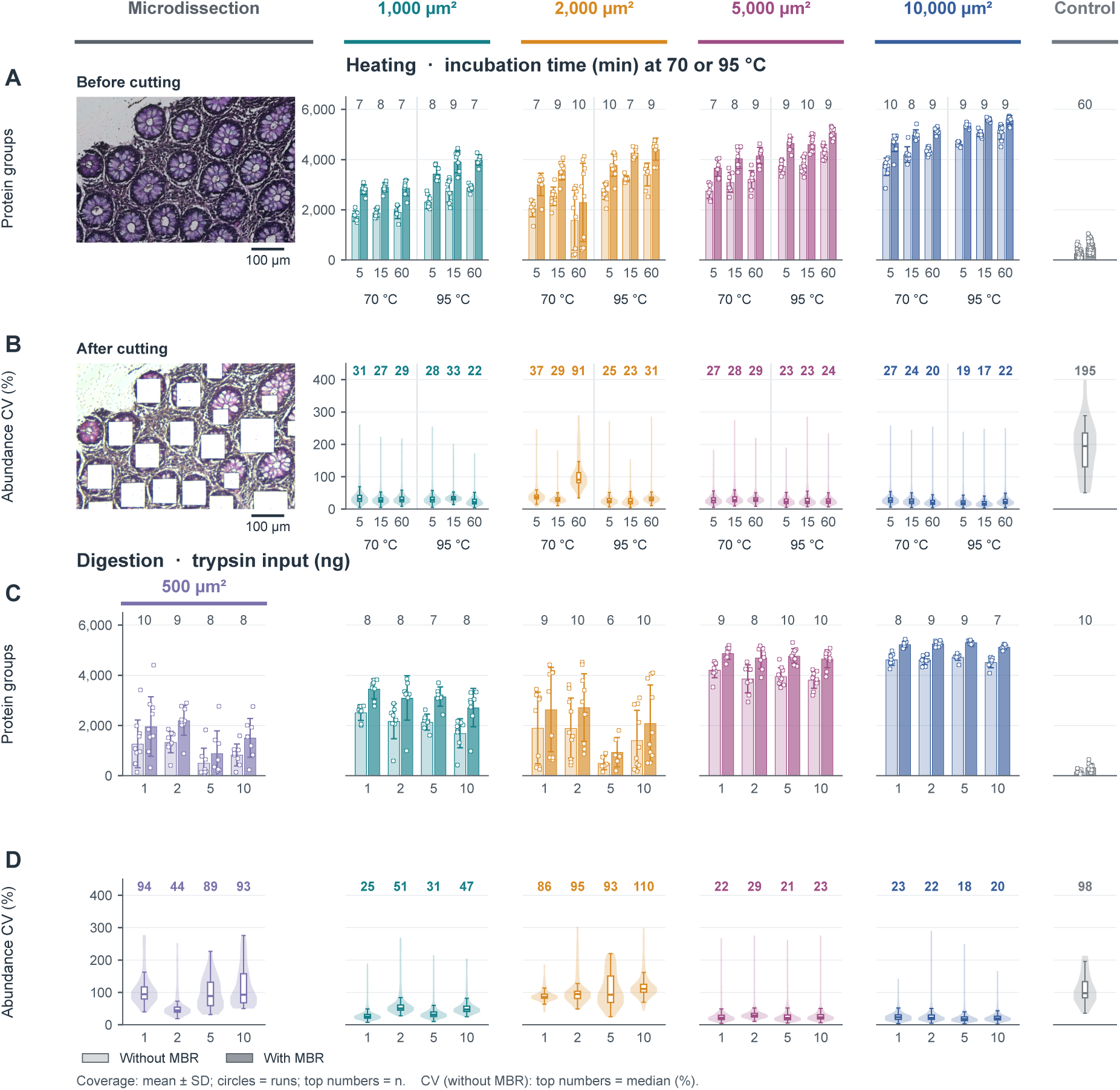
Optimizing sample preparation. (A,B) Protein-group coverage and abundance coefficients of variation (CVs) after heating at 70 or 95 °C for 5, 15, or 60 minutes. (C,D) Corresponding measurements after digestion with 1, 2, 5, or 10 ng trypsin. Colors identify tissue inputs. Lighter and darker bars show mean coverage ± SD without and with MBR, respectively, for the same retained runs. Circles show individual runs, and numbers above bar pairs indicate sample counts. Panels B and D show CVs without MBR on a shared linear scale, with median CVs above the violins. Each tested condition contained ten measurements per tissue input before QC, with 500 µm² tested only in the digestion experiment. The images show tissue before and after collection. Scale bars, 100 µm.

We next evaluated the digestion step by comparing 1, 2, 5, and 10 ng trypsin across 500–10,000 µm² tissue inputs (Figure 2C). Quality control excluded 29 of 210 measurements. With 1 ng trypsin, mean coverage ranged from 1,268 protein groups at 500 µm² to 4,625 at 10,000 µm². Doubling the amount to 2 ng produced changes ranging from 14% lower to 5% higher coverage across tissue inputs, and further increases to 5 or 10 ng provided no consistent benefit. At 5,000 and 10,000 µm², mean coverage with 2–10 ng remained within 10% of that obtained with 1 ng, while at 500–2,000 µm², 10 ng yielded 26–35% fewer protein groups than 1 ng. The 2,000 µm² condition with 5 ng trypsin yielded particularly low coverage (505 ± 269 protein groups; mean ± SD), 73% below that obtained with 1 ng. Median abundance CVs were approximately 18–29% at 5,000 and 10,000 µm², compared with 44–94% at 500 µm² and 86–110% at 2,000 µm² (Figure 2D).

With MBR, mean coverage increased across all heating and digestion conditions. At 95 °C for 15 minutes, mean counts were 3,916, 4,275, 4,606, and 5,600 protein groups at 1,000, 2,000, 5,000, and 10,000 µm², respectively.

### Assessing the analytical performance of the workflow

Using the selected 95 °C/15-minute heating condition, we next assessed how tissue input affected the analytical performance of the workflow across 1,000, 2,000, 5,000, and 10,000 µm² regions. Quality control retained 35 of 40 tissue measurements. Reducing tissue area from 10,000 to 1,000 µm² decreased mean precursor coverage from 28,903 to 12,546, a reduction of 57% (Figure 3A).

**Figure 3.**
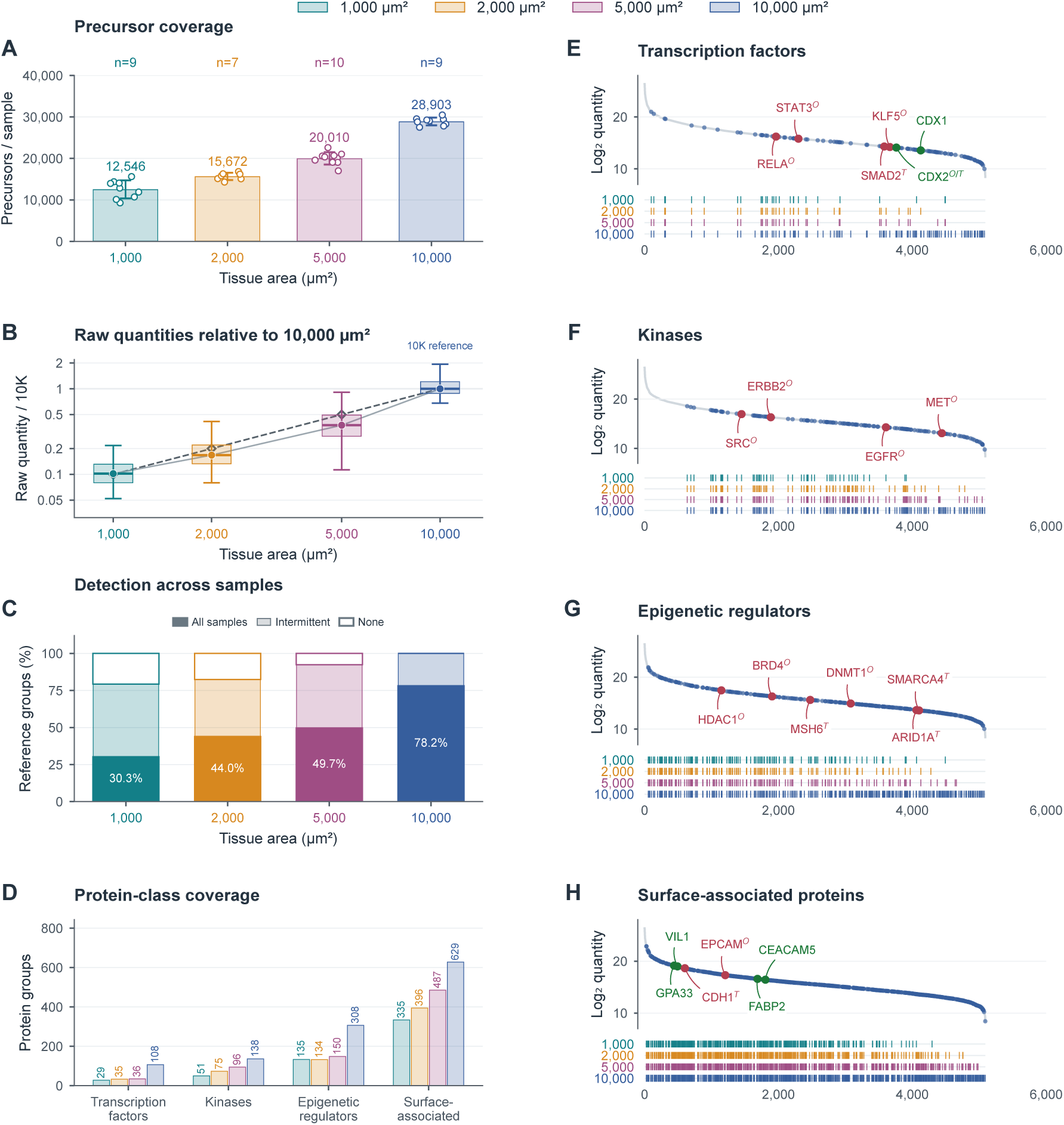
Quality assessment of the workflow. (A) Number of precursors identified per retained sample after heating at 95 °C for 15 minutes, shown as mean ± SD with individual measurements and sample counts. (B) Raw quantities relative to the 10,000 µm² reference, using 4,466 precursors from 1,308 protein groups quantified in all 35 retained measurements. The dashed line shows the area-proportional expectation; quantities were not normalized across samples. (C) Detection within the 5,090-group reference core, distinguishing groups found in every sample (filled), intermittently (pale), or in none (white). (D) Protein-class counts for groups detected in at least half the retained measurements at each input. From 1,000 to 10,000 µm², these counts increased by 272% for transcription factors, 171% for kinases, 128% for epigenetic regulators, and 88% for surface-associated proteins. (E–H) Abundance-ranked profiles at 10,000 µm², with the reference proteome in grey and members of each protein class in blue. Colored tracks indicate consistent detection at each tissue input. Green labels identify intestinal markers and red labels identify cancer-associated genes; O, T, and O/T denote oncogene, tumour-suppressor, and dual annotations. All data were searched without MBR.

To examine whether quantities scaled with tissue area, we compared the same 4,466 precursors quantified in all 35 retained measurements. Median relative quantities at 1,000, 2,000, and 5,000 µm² were 10%, 17%, and 38% of the 10,000 µm² reference, compared with area-proportional expectations of 10%, 20%, and 50% (Figure 3B). Quantities therefore broadly followed the expected reduction in sampled material, closely matching the proportional expectation at 1,000 µm² and falling below it at the intermediate inputs. Detection also became more intermittent at smaller inputs, as expected. The fraction of the reference proteome detected in every retained measurement fell from 78% at 10,000 µm² to 30% at 1,000 µm², while the fraction detected in only some measurements rose from 22 to 49% (Figure 3C). Greater sample-to-sample heterogeneity in smaller tissue regions, together with lower quantities approaching the detection limit, may contribute to this pattern.

To assess the biological clinical relevance of the coverage retained at smaller inputs, we focused on four protein classes that include drug targets and biomarkers: epigenetic regulators, kinases, transcription factors, and surface-associated proteins. Members of all four classes remained detectable at 1,000 µm² (Figure 3D), retaining 27–52% of the corresponding 10,000 µm² reference classes, with the lowest retention for transcription factors and the highest for surface-associated proteins. The abundance-ranked profiles showed that the loss of coverage mainly affected lower-abundance proteins (Figure 3E– H).

The proteins detected at low input included markers of the intestinal epithelium. Villin-1 (VIL1), CEACAM5, and GPA33 were detected in every retained measurement across all four areas. At 1,000 µm², FABP2 and ANPEP were detected in seven of nine and four of nine measurements, respectively; both were detected in every retained measurement at larger inputs. The profiles also contained proteins encoded by genes annotated in OncoKB as oncogenes (EGFR, ERBB2, SRC, and BRD4) or tumour suppressors (CDH1 and SMARCA4) (Figure 3E–H).

### Assessing tissue collection in a 1,536-well plate

To further increase sample throughput and facilitate greater automation of the spatial proteomics workflow, we investigated the feasibility of collecting microdissected tissue directly into 1,536-well plates. In addition to increasing the number of samples that could be processed within a single plate, we sought to assess whether tissue could be reliably collected into the intended wells at this higher plate density (and smaller collection well). To evaluate this, we collected material into a 1,536-well plate, with ten target wells each for 1,000, 2,500, 5,000, 7,500, and 10,000 µm² tissue inputs. Direct injection from this format was not possible with the Vanquish Neo setup used here, so samples were transferred to 384-well plates before LC–MS analysis. This additional handling was expected to reduce coverage through sample losses during transfer.

Proteomic detections were mapped back to 357 reported source wells, comprising 50 intended collection wells and 307 surrounding wells (Figure 4). The strongest detections generally coincided with the intended wells, supporting collection at the assigned positions. However, some wells indicated potential collection errors. For example, the intended well S6 yielded few detections, whereas the adjacent untargeted well S7 yielded substantially more.

**Figure 4.**
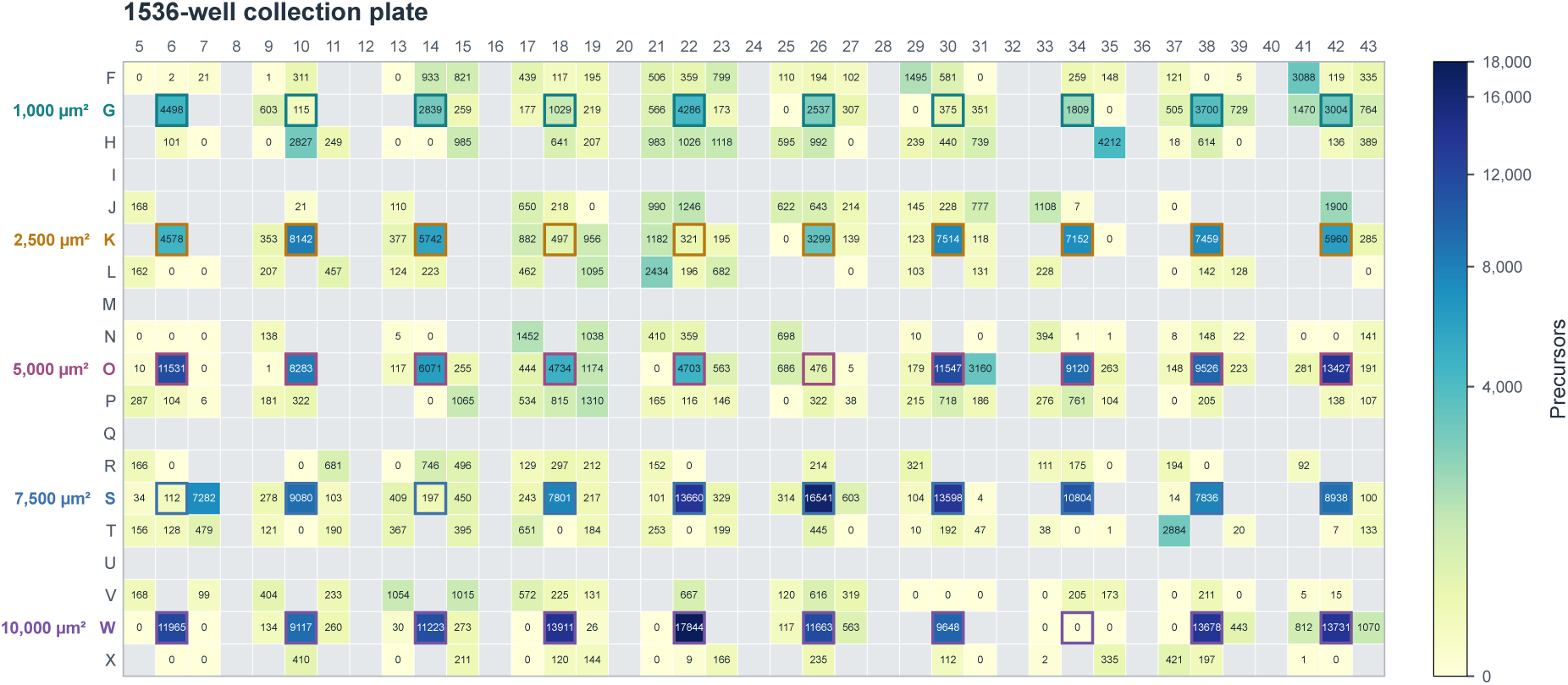
Assessment of tissue placement in a 1,536-well plate. Distribution of proteomic detections across intended collection wells and surrounding wells, shown as the number of unique quality-filtered precursors after transfer to 384-well plates for LC–MS analysis. The map shows the reported region at its original plate coordinates. Gray cells indicate wells with no reported run. Colored outlines mark the 50 tissue targets, with ten wells each for 1,000, 2,500, 5,000, 7,500, and 10,000 µm² in rows G, K, O, S, and W, respectively. Numbers indicate precursor counts, while the color scale uses square-root spacing to show variation in wells with lower coverage.

### ShapeX connects image annotations with one-pot sample preparation

Alongside the preparation workflow, we developed ShapeX as an integrated QuPath extension to make the transition from tissue annotation to sample collection and preparation more accessible (Figure 5). ShapeX works with annotations already present in a QuPath project, allowing regions to be defined using manual annotation or the image-analysis tools available within QuPath. The extension then connects these regions to a collection plan, rather than requiring users to manage image coordinates, well assignments, and preparation files independently.

**Figure 5.**
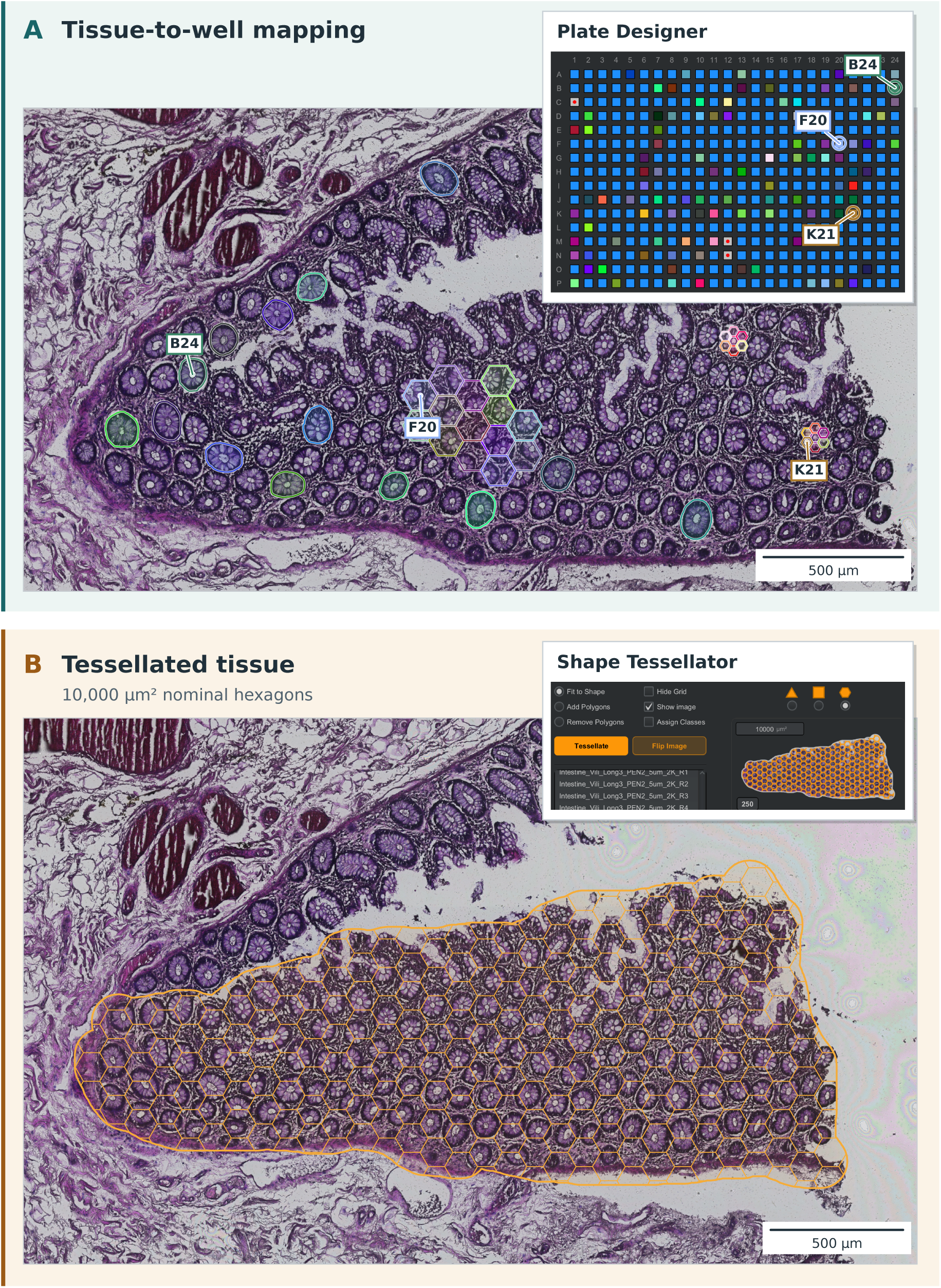
ShapeX links tissue regions with plate allocation and spatial sampling. (A) Matching labels link selected tissue annotations to their assigned collection wells in the Plate Designer inset. (B) The Shape Tessellator inset previews the division of a selected region into hexagons with a nominal area of 10,000 µm², shown overlaid on the tissue.

The Shape Tessellator divides selected annotations into triangular, square, or hexagonal sampling regions with a user-defined area. Previewing the grid over the image allows users to examine the proposed sampling scale before creating the annotations. The resulting regions remain QuPath annotations, preserving their relationship to the source image and providing the spatial information needed for later reconstruction. Tessellation therefore offers a practical way to vary the sampled area in experiments of the type examined here.

The Plate Designer supports standard 96-, 384-, and 1,536-well formats, as well as custom row and column dimensions. Annotations can be allocated individually, pooled according to their class, or organized so that each slide is assigned to its own plate. Users can inspect and adjust the assignments visually, reserve unavailable wells, and include negative controls. Sample randomization includes an explicit seed, allowing an allocation to be reproduced. These options connect experimental decisions, such as separating tissue classes or distributing samples across a plate, with the actual collection layout.

For laser microdissection, ShapeX exports annotation boundaries and collection-well assignments in Leica LMD XML format, including the three calibration points needed for transfer to the dissection system. Plate overview files record the sampling layout and associated sample information in spreadsheet or text formats. Annotation geometry, sample metadata, and well locations can additionally be exported in AnnData format for integration with proteomic measurements and downstream analysis with Scanpy and other tools in the scverse ecosystem [17, 18]. These exports preserve the link between the original tissue region and its destination well.

### ShapeX reconstructs spatial protein distributions in intestinal tissue

To evaluate spatial reconstruction, we linked proteomic measurements from two intestinal experiments to their original tissue annotations (Figure 6). In the longitudinal experiment, regions were sampled along both sides of intestinal structures. L1 and R1 marked the base, and increasing segment numbers followed each side toward the luminal end, labelled as the tip (Figure 6A). We pooled observed abundance values from all collected villi by side and segment number and mapped the medians onto a representative tissue outline (Figure 6B,C).

**Figure 6.**
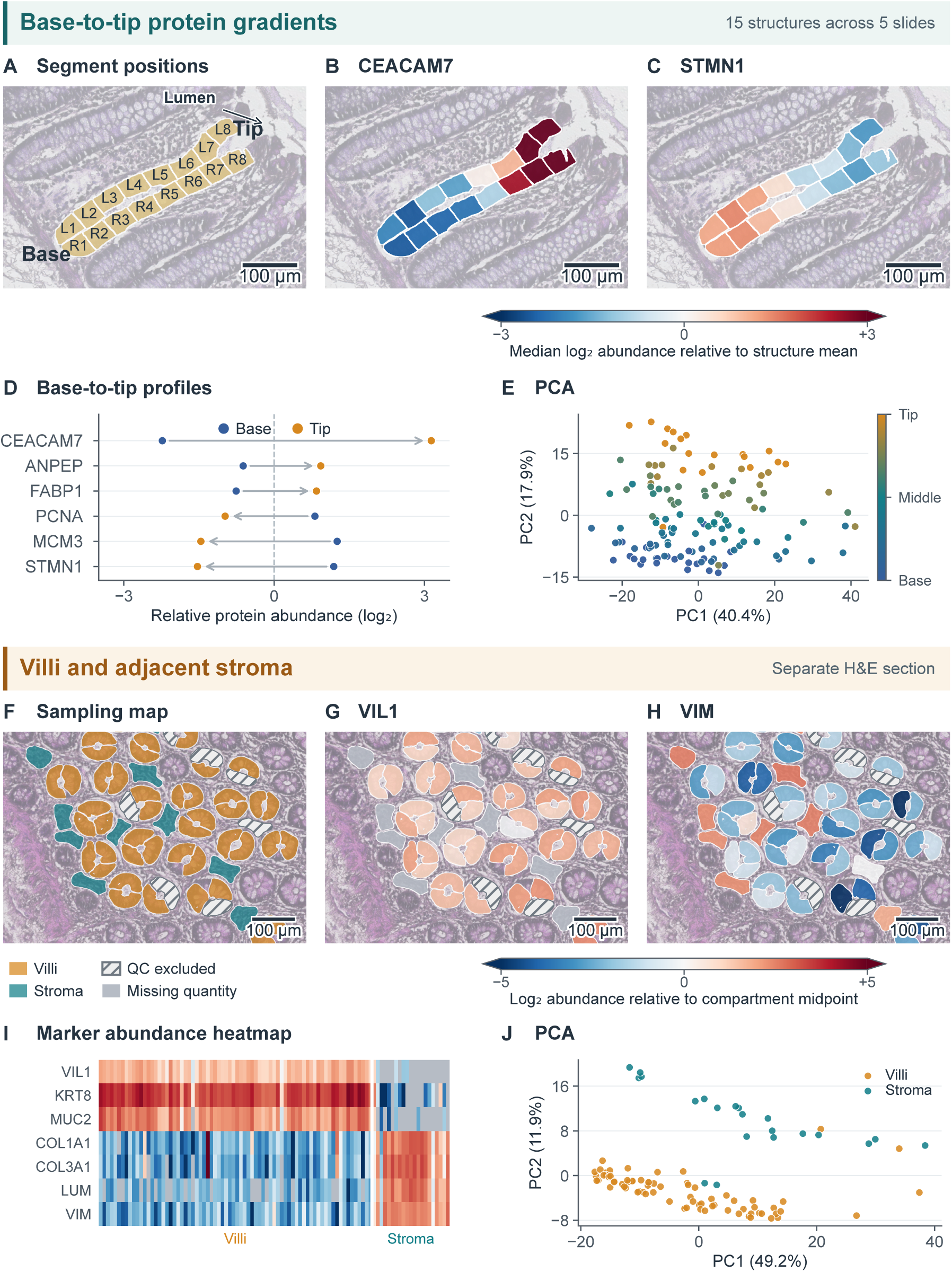
ShapeX reconstructs protein gradients along intestinal structures and contrasts between villi and stroma. (A) Segment positions on a arbitrarily selected longitudinal villi. (B,C) Median CEACAM7 and STMN1 abundances at each side and segment number, calculated from observed values across all collected villi and mapped onto the selected villi. (D) Median base and tip abundances across structures with observed quantities in both zones. In B–D, each protein’s log_2_ abundances are centered on its mean within each sampled structure. (E) PCA colored by normalized base-to-tip position. (F–H) Sampling map and VIL1 and VIM abundances in a separate HCE section. (I) Heatmap of seven markers, grouped by compartment and ordered by position on the slide. Grey indicates missing quantities. In G–I, log_2_ abundances are relative to the midpoint of each protein’s villi and stroma medians. (J) PCA of median-normalized log_2_ abundances for protein groups quantified in all analyzed regions, colored by compartment. Scale bars, 100 µm.

CEACAM7 was enriched toward the lumen, whereas STMN1 was more abundant near the base (Figure 6B–D). ANPEP and FABP1 also increased toward the luminal end, while PCNA and MCM3 showed higher abundance at the base (Figure 6D). The CEACAM7 pattern agrees with its reported localization in differentiated epithelial cells of the upper colonic crypt [19]. The luminal trends of ANPEP and FABP1 agree with previously reported intestinal expression patterns [20], and the basal enrichment of PCNA and MCM3 is consistent with their roles in DNA replication [21, 22]. Position along the sampled axis was also reflected in the broader proteome. PC2 explained 17.9% of the variance and tracked normalized base-to-tip position (Spearman *ρ* = 0.903, Figure 6E).

In a separate HCE section, we compared sampled glandular epithelium (villi) with the adjacent interglandular stroma (Figure 6F–J). Among observed quantities, median VIL1 abundance was 3.11 log_2_ units higher in villi, while median VIM abundance was 4.67 log_2_ units higher in stroma. Mapping the two proteins back to the tissue showed how these contrasts corresponded to the annotated compartments (Figure 6F–H). The heatmap also showed higher abundances of KRT8 and MUC2 in villi and of COL1A1, COL3A1, and LUM in stroma (Figure 6I). These patterns agree with the reported distribution of epithelial and stromal markers in colorectal tissue [23, 24]. PCA of the compartment profiles separated most villi and stroma regions along PC2, which explained 11.9% of the variance (Figure 6J).

## Discussion

The one-pot workflow connects LCM tissue collection with proteomic analysis by retaining samples in the same well through heating, digestion, and direct LC–MS aspiration. The optimization experiments support 95 °C heating and low trypsin input as practical preparation conditions. Heating at 95 °C generally yielded greater coverage than heating at 70 °C. Without MBR, extending incubation from 15 to 60 minutes provided modest gains of approximately 1–5% at most tissue inputs. The longer incubation may therefore be useful when proteome depth is prioritized, but its additional processing time reduces sample-preparation throughput. A 15-minute incubation offers a practical balance between coverage and processing time. Increasing trypsin above 1 ng yielded no consistent gain in coverage. Reduced coverage at some higher inputs may reflect signal suppression from trypsin-derived peptides. The searches with MBR showed similar preparation trends, with the greatest relative gains in coverage at smaller tissue inputs.

The intestinal experiments demonstrate how ShapeX can link proteomic measurements to tissue morphology. Along the sampled structures, the opposing gradients are consistent with differentiated epithelial functions toward the lumen and proliferation near the base. The directions persisted across filtering and normalization sensitivity analyses. In the separate HCE section, mapping VIL1 and VIM back to the annotations showed abundance differences between glandular epithelium and adjacent stroma. These examples illustrate how the same annotation-to-well workflow can support both longitudinal sampling and comparisons between neighboring compartments.

Collection into 1,536-well plates is a viable strategy for future scaling. The strongest detections generally coincided with the intended collection wells, but substantial signal in some surrounding wells highlights the need to improve collection precision and assess cross-sample contamination. Further tuning of plate alignment and collection geometry, together with checks of material recovered in neighboring wells, will be necessary to establish reliable collection and prevent cross-sample contamination at this density. The present experiment establishes the feasibility of high-density collection, although the LC setup required transfer to 384-well plates. Direct injection from 1,536-well plates could substantially increase proteome depth but remains to be evaluated.

Throughout all experiments, we observed no abnormal column degradation or other LC– MS problems attributable to direct injection of the prepared digests, despite residual tissue material in the wells. These observations support a desalting- and clean-up-free preparation strategy without requiring an Evosep system or transfer to Evotips. Needle access also depends on the plate seal. We used pierceable foil seals in the preparation and collection-plate experiments to support direct aspiration.

ShapeX complements open-source frameworks, such as openDVP [25], that uses the QuPath-to-LMD web application to convert exported QuPath annotations into microdissection files. As a QuPath extension, ShapeX works directly with annotations in the project, removing the intermediate annotation export. Users can tessellate regions, design collection plates, and randomize well assignments within QuPath. ShapeX then generates Leica LMD and reagent-dispensing files from the same plate layout, preserving the link between each sampled region and its collection well for spatial reconstruction. Combined with one-pot sample preparation, this integration could make spatial proteomics more accessible and support profiling of larger clinical cohorts to investigate how molecular changes relate to tissue pathology.

## Methods

### Tissue preparation

All experiments used tissue from the same formalin-fixed, paraffin-embedded (FFPE) human colon block purchased from Avaden BioSciences Inc. (Seattle, WA, USA). The supplied specimen record described a resection collected on 20 October 2015 from a 29-year-old female. According to the supplier’s ethics statement dated 13 March 2019, human samples were collected from US hospitals under informed consent compliant with the Health Insurance Portability and Accountability Act (HIPAA). Consent forms and collection protocols were approved by independent review boards at the supplying hospitals, and the consent included commercial research use.

The supplier reports standardized collection, processing, and storage procedures within US facilities accredited by the College of American Pathologists (CAP) and certified under the Clinical Laboratory Improvement Amendments (CLIA). Its banked-specimen workflow includes pathology review and final HCE imaging and quality control. Before staining, 5 µm sections were mounted on 1 mm glass slides bearing a 2.0 µm polyethylene naphthalate (PEN) membrane (MembraneSlide 1.0 PEN, order no. 415190-9042-000, Carl Zeiss Microscopy GmbH, Jena, Germany).

### Hematoxylin–eosin (HsE) staining

HCE staining followed the previously described protocol [16]. After adhesion, sections were deparaffinized in Tissue-Tek Tissue-Clear Xylene Substitute for 10 min, followed by 20 dips in 99% ethanol, 10 dips in 96% ethanol, and 10 dips in 70% ethanol. Slides were rinsed in running demineralized water for 5 min and stained with Mayer’s hematoxylin for 4 min. After a 5-minute rinse in running water, sections were stained for 1 min with 0.1% eosin in 0.1 M Walpole’s acetate buffer, followed by a final 3-minute rinse in running water. Slides were left to dry overnight in a fume hood and then stored in a closed container at room temperature.

### Image acquisition and LCM collection

After HCE staining, PEN membrane slides were scanned in brightfield using a NanoZoomer XR digital slide scanner (Hamamatsu) at 40× magnification. To improve image quality, holes were punctured in the PEN membrane using a ZEISS PALM MicroBeam prior to imaging. Whole-slide images were acquired in NDPI format using NDP.scan (v3.4.3), with a calibrated pixel size of approximately 0.23 µm/pixel, and imported into QuPath (v0.7.0) for annotation and subsequent sample-to-well mapping and export using ShapeX.

Regions of interest were annotated in QuPath and assigned to collection wells using ShapeX. Annotation boundaries, well assignments, and three calibration points surrounding the selected regions were exported in Leica LMD XML format. The corresponding files were imported into the Leica LMD6 control software (v8.5.0.91.36), and the calibration points were relocated on each membrane slide before dissection. ShapeX also exported plate layouts and dispensing files for automated sample preparation, together with AnnData files retaining sample identities, annotation geometry, and collection-well assignments for spatial reconstruction.

Tissue was collected at 40× magnification using laser power 16, aperture 1, speed 16, middle pulse count 1, and final pulse 10. Dissected regions were collected directly into Eppendorf twin.tec PCR Plate 384 LoBind plates (Cat. No. 0030129547). A custom 3D-printed adapter raised the collection plate by 6.6 mm toward the tissue slide; the same adapter was used for the 1,536-well collection experiment.

For the heating experiment, regions of 1,000, 2,000, 5,000, and 10,000 µm² were collected alongside empty-well controls. The digestion experiment additionally included 500 µm² regions. For longitudinal intestinal sampling, 15 structures across five PEN slides were divided into consecutive regions of approximately 2,000 µm² along their two sides. Segment identifiers retained the slide, structure, side, and position, with L1 and R1 marking the base and increasing segment numbers following the structure toward the luminal end.

Collection into the 1,536-well format was evaluated using polypropylene MASTERBLOCK deep-well plates with V-bottom wells (Greiner Bio-One, Cat. No. 782261; 32 rows and 48 columns). Ten target wells were assigned to each of five tissue inputs: 1,000, 2,500, 5,000, 7,500, and 10,000 µm². Targets occupied rows G, K, O, S, and W, respectively, at four-column intervals from column 6 to column 42. Surrounding wells were analyzed to assess tissue placement. Samples from this collection plate were transferred to two 384-well plates before LC–MS/MS analysis because direct aspiration from the 1,536-well format was not possible with the Vanquish Neo configuration used here. A separate 384-well collection control comprised 50 tissue samples and nine empty controls.

### Sample processing

Samples were prepared in 384-well plates, with heating conditions and trypsin input varied in the optimization experiments. After collection, acetonitrile (50 µL per well; LiChrosolv Hypergrade for LC–MS, Supelco, Cat. No. 1000292500) was dispensed using an Opentrons Flex, and the plates were dried for 60 min in a Savant SpeedVac SPD120 vacuum concentrator. Dried plates were covered and stored at −80 °C until further processing.

Plates were thawed at room temperature and centrifuged at 4,347 × g for 1 min. An I.DOT S dispenser (Dispendix) was used to add 1 µL of lysis buffer containing 100 mM triethylammonium bicarbonate (TEAB; Thermo Scientific, Cat. No. 90114) and 0.01% n-dodecyl-β-D-maltoside (DDM; Cat. No. D4641-500MG) in LC–MS-grade water (LiChrosolv, Supelco, Cat. No. 1153332500). Plates were centrifuged for 1 min at 4,347 × g and heated in a Biometra TAdvanced thermal cycler. The standard incubation was 5 min at 95 °C. The heating experiment compared 70 and 95 °C for 5, 15, and 60 min across the four tissue inputs, with matched empty-well controls. The 95 °C/15-minute condition was used for the subsequent tissue-area performance analysis.

### Digestion and sample storage

Trypsin Platinum, Mass Spec Grade (Promega), was dispensed using the I.DOT S. The standard protocol used 1 µL of 10 ng/µL trypsin per well. In the digestion experiment, the total trypsin input was varied between 1, 2, 5, and 10 ng across tissue areas of 500, 1,000, 2,000, 5,000, and 10,000 µm², with ten measurements per combination and ten empty-well controls at 10 ng trypsin. After enzyme addition, plates were centrifuged at 4,347 × g for 1 min and incubated at 37 °C for 18 h. Digestion was quenched by adding 1 µL of 1% trifluoroacetic acid (TFA; Sigma-Aldrich, Cat. No. 302031-10X1ML). Plates were centrifuged again for 1 min at 4,347 × g and stored at −80 °C until analysis. Reagent concentrations refer to the dispensed solutions.

Longitudinal intestinal samples were injected through silicone microplate lids. For the preparation and 1,536-well collection experiments, plates were sealed with Pierceable Foil Heat Seal (Bio-Rad, Cat. No. 1814040) using a PX1 PCR Plate Sealer at 170 °C for 3 s, and the autosampler needle pierced the foil directly for aspiration.

### Liquid chromatography settings

#### Preparation and collection-plate experiments

Peptides were separated on a Vanquish Neo UHPLC system using a PepMap Neo C18 trap cartridge (5 mm × 300 µm internal diameter, 5 µm particles, Cat. No. 174500, Thermo Fisher Scientific) and an Aurora Rapid XT C18 analytical column (8 cm × 75 µm internal diameter, 1.7 µm particles, Cat. No. AUR5-8075C18-XT, IonOpticks).

The 8.5-minute LC program started at 8% B and 800 nL/min. Buffer B increased to 10% at 0.075 min, 18% at 0.263 min, and 35% at 1.451 min. From 1.451 to 1.501 min, buffer B increased to 99% and the flow rate decreased to 130 nL/min. The column was washed at these settings until 8.450 min. Buffer B and the flow rate then returned to 4% and 800 nL/min, respectively, by 8.500 min for column equilibration.

#### Longitudinal intestinal samples

Peptides were separated using a Vanquish Neo UHPLC system connected to a 50 cm C18 OptiSpray µPAC Neo cartridge with an 8 µm pulled emitter (Thermo Fisher Scientific). The column was maintained at 50 °C. The method used a 12.5-minute gradient and 2.5 min of loading overhead, corresponding to 96 samples per day. The spray voltage was 1.8 kV.

Buffer B consisted of 80% acetonitrile in water with 0.1% formic acid. Buffer B increased from 1 to 10% over 0–0.05 min, from 10 to 24.5% over 0.5–1.95 min, and from 24.5 to 37.5% over 1.95–3.2 min at a nominal flow rate of 750 nL/min. The flow rate was then reduced to 250 nL/min while buffer B increased from 37.5 to 40% over 3.2–3.3 min and from 40 to 90% over 3.3–3.4 min, followed by a 9.1-minute hold.

### Mass spectrometry settings

#### Preparation and collection-plate experiments

Data were acquired on an Orbitrap Astral mass spectrometer equipped with an EASY-Spray source and a FAIMS interface (Thermo Fisher Scientific). The instrument operated in positive-ion mode at a spray voltage of 1.6 kV and an ion transfer tube temperature of 240 °C. The FAIMS compensation voltage was −48 V, with inner and outer electrode temperatures of 100 and 80 °C, respectively, and a carrier gas flow of 3.6 L/min.

Orbitrap MS1 spectra were acquired at a resolution of 240,000 over 400–800 m/z, with a normalized automatic gain control (AGC) target of 500% and a maximum injection time of 100 ms. Data-independent acquisition (DIA) used 67 non-overlapping isolation windows of 6 m/z across the precursor range of 400–800 m/z. Fragment ions were recorded in the Astral analyzer over 150–2,000 m/z, with a normalized AGC target of 500% and a maximum injection time of 10 ms. Precursors were fragmented by higher-energy collisional dissociation at a normalized collision energy of 25%, with N control set to 43 spectra. The radiofrequency lens was set to 45%, and spectra were recorded in profile mode with one microscan. The expected LC peak width was 6 s and the default charge state was 2.

#### Longitudinal intestinal samples

Data were acquired on an Orbitrap Astral mass spectrometer equipped with a FAIMS Pro interface (Thermo Fisher Scientific). The instrument operated in positive-ion mode at a FAIMS compensation voltage of −45 V, with inner and outer electrode temperatures of 100 and 80 °C, respectively. Orbitrap MS1 spectra were acquired at a resolution of 240,000 over 400–800 m/z, with a normalized automatic gain control (AGC) target of 500% and a maximum injection time of 100 ms. Data-independent acquisition (DIA) MS2 spectra were acquired in the Astral analyzer over 150–2,000 m/z using 4 m/z isolation windows with 1 m/z overlap. N control cycled every 50 scans. The MS2 AGC target was 500%, and the maximum injection time was 10 ms. Precursor ions were fragmented by higher-energy collisional dissociation at a normalized collision energy of 25%.

### Data analysis

All raw MS data were searched in DIA-NN (v2.7.0) [26] using a spectral library generated from reviewed human UniProt sequences, including isoforms (downloaded 9 November 2026). Library generation and searches used Trypsin/P with one missed cleavage. Standard settings were used with fixed cysteine carbamidomethylation disabled and methionine oxidation and protein N-terminal acetylation included as variable modifications. The heating and digestion datasets were searched with and without match between runs (MBR). MBR was also enabled for the villi-versus-stroma and longitudinal intestinal datasets. All other search settings were identical across experiments. For the coverage comparison in Figure 2, the same filters were applied to both reports, and counts were summarized over the runs retained by coverage-based quality control of the searches without MBR.

Follow-up data analysis was performed in Python using several packages. pandas and NumPy were used for data wrangling, Matplotlib for visualization, SciPy for statistical analysis, and scikit-learn for principal component analysis. AnnData and Shapely were used to link proteomic measurements to the original ShapeX annotations and reconstruct spatial protein distributions (a guide to installing ShapeX is available on GitHub: https://github.com/Schoof-Lab/ShapeX).

## List of abbreviations

3D: Three-dimensional
AGC: Automatic gain control
AI: Artificial intelligence
ANPEP: Alanyl aminopeptidase, membrane
ARID1A: AT-rich interactive domain-containing protein 1A
BRD4: Bromodomain-containing protein 4
CAP: College of American Pathologists
CDH1: Cadherin-1
CDX1: Homeobox protein CDX-1
CDX2: Homeobox protein CDX-2
CEACAM5: Carcinoembryonic antigen-related cell adhesion molecule 5
CEACAM7: Carcinoembryonic antigen-related cell adhesion molecule 7
CLIA: Clinical Laboratory Improvement Amendments
COL1A1: Collagen alpha-1(I) chain
COL3A1: Collagen alpha-1(III) chain
CV: Coefficient of variation
DDM: n-Dodecyl-β-D-maltoside
DIA: Data-independent acquisition
DIA-NN: Data-independent acquisition–neural networks
DNA: Deoxyribonucleic acid
DNMT1: DNA (cytosine-5)-methyltransferase 1
EGFR: Epidermal growth factor receptor
EPCAM: Epithelial cell adhesion molecule
ERBB2: Erb-b2 receptor tyrosine kinase 2
FABP1: Fatty acid-binding protein 1
FABP2: Fatty acid-binding protein 2
FAIMS: High-field asymmetric waveform ion mobility spectrometry
FASP: Filter-aided sample preparation
FFPE: Formalin-fixed, paraffin-embedded
GPA33: Glycoprotein A33
HsE: Hematoxylin and eosin
HDAC1: Histone deacetylase 1
HIPAA: Health Insurance Portability and Accountability Act
KLF5: Krueppel-like factor 5
KRT8: Keratin, type II cytoskeletal 8
LC: Liquid chromatography
LC–MS: Liquid chromatography–mass spectrometry
LC–MS/MS: Liquid chromatography–tandem mass spectrometry
LCM / LMD: Laser capture microdissection / laser microdissection
LUM: Lumican
MALDI-MSI: Matrix-assisted laser desorption/ionization mass spectrometry imaging
MBR: Match between runs
MCM3: Minichromosome maintenance complex component 3
MET: Hepatocyte growth factor receptor
MMI: Molecular Machines & Industries
MS: Mass spectrometry
MS1 / MS2: Precursor-ion / fragment-ion mass spectra
MSH6: DNA mismatch repair protein Msh6
MSI: Mass spectrometry imaging
MUC2: Mucin-2
m/z: Mass-to-charge ratio
NDPI: NanoZoomer digital slide image format
OncoKB: Oncology Knowledge Base
openDVP: Open deep visual proteomics
PC1 / PC2: First / second principal component
PCA: Principal component analysis
PCNA: Proliferating cell nuclear antigen
PCR: Polymerase chain reaction
PEN: Polyethylene naphthalate
QC: Quality control
RELA: Transcription factor p65
SD: Standard deviation
SMAD2: SMAD family member 2
SMARCA4: SWI/SNF-related BAF chromatin remodeling complex subunit ATPase 4
SP3: Single-pot, solid-phase-enhanced sample preparation
SRC: SRC proto-oncogene, non-receptor tyrosine kinase
STAT3: Signal transducer and activator of transcription 3
STMN1: Stathmin-1
TEAB: Triethylammonium bicarbonate
TFA: Trifluoroacetic acid
UHPLC: Ultra-high-performance liquid chromatography
µPAC: Micropillar array column
VIL1: Villin-1
VIM: Vimentin
XML: Extensible Markup Language

## Declarations

### Ethics approval

Human FFPE colon tissue was purchased from Avaden BioSciences Inc. According to the supplier’s ethics statement, specimens were collected under HIPAA-compliant informed consent, with consent forms and collection protocols approved by independent review boards at the supplying US hospitals.

### Author contributions

R.D., V.P., and E.M.S conceived and designed the project. R.D. and M.W.N. performed the experimental investigations, while J.B.J. and V.V.T. aided in workflow optimization. Sectioning, staining, and imaging of tissue sections were conducted by R.D., N.M.H., and J.B.J. ShapeX was developed by R.D. with the aid of an LLM. Data analysis was performed by R.D. The original manuscript was drafted by R.D., with all other authors contributing to its review and editing. V.P. and E.M.S. supervised the work.

## Acknowledgements

We thank all members of the Cell Diversity Lab, for their constructive input and valuable discussions.

